# Emetine as an antiviral agent suppresses SARS-CoV-2 replication by inhibitinginteraction of viral mRNAwith eIF4E: An *in vitro* study

**DOI:** 10.1101/2020.11.29.401984

**Authors:** Ram Kumar, Nitin Khandelwal, Yogesh Chander, Thachamvally Riyesh, Baldev R. Gulati, Yash Pal, Bhupendra N. Tripathi, Sanjay Barua, Naveen Kumar

## Abstract

Emetine is a FDA-approved drug for the treatment of amebiasis. In the recent times we had also demonstrated the antiviral efficacy of emetine against some RNA and DNA viruses. Following emergence of the COVID-19, we further evaluated the*in vitro* antiviral activity of emetine against severe acute respiratory syndrome coronavirus 2 (SARS-CoV-2). The therapeutic index of emetine was determined to be 10910.4, at a cytotoxic concentration 50 (CC_50_) of 1603.8 nM and effective concentration 50 (EC_50_) of 0.147 nM.Besides, we also demonstrated the protective efficacy of emetine against lethal challenge with infectious bronchitis virus (IBV; a chicken coronavirus) in the embryonated chicken egg infection model. Emetine treatment was shown to decrease viral RNA and protein synthesis without affecting other steps of viral life cycle such as attachment, entry and budding.In a chromatin immunoprecipitation (CHIP) assay, emetine was shown to disrupt the binding of SARS-CoV-2 RNA with eIF4E (eukaryotic translation initiation factor 4E, a cellular cap-binding protein required for initiation ofprotein translation). Further, SARS-CoV-2 was shown to exploit ERK/MNK1/eIF4E signalling pathwayfor its effective replication in the target cells. To conclude, emetine targets SARS-CoV-2 protein synthesis which is mediated via inhibiting the interaction of SARS-CoV-2 RNA with eIF4E. This is a novel mechanistic insight on the antiviral efficacy of emetine. *In vitro* antiviral efficacy against SARS-CoV-2 and its ability to protect chicken embryos against IBV suggests that emetine could be repurposed to treat COVID-19.

## 1. Introduction

The ongoing pandemic of COVID-19 has become a health emergency of international concern. Therapeutic agents that can be used against COVID-19 or other coronavirus diseases are unavailable.Emetine dihydrochloride hydrate or emetine isa natural alkaloid found in three plant families:*Alangiaceae, Icacinaceae, and Rubiaceae*. Emetine is a FDA-approved drug for the treatment of amebiasis(Bleasel and Peterson, 2020).

Emetine has beenshownto disrupt protein synthesis in mammalian, yeast and plant cells (Grollman, 1966, 1968; Gupta and Siminovitch, 1976; Han et al., 2014; Jimenez et al., 1977; Smirnova et al., 2003). Recent studies by our and other groups have suggested *in vitro*antiviral efficacy of emetine against some RNA and DNA viruses (Chaves Valadao et al., 2015; Choy et al., 2020; Deng et al., 2007; Khandelwal et al., 2017; Mukhopadhyay et al., 2016; Ramabhadran and Thach, 1980; Yang et al., 2018).Emetine can directly inhibit viral polymerase (Chaves Valadao et al., 2015; Khandelwal et al., 2017; Yang et al., 2018), although the major antiviral activity of emetine is believed to be mediated via targeting certain cellular factors (Khandelwal et al., 2017). While most studies are based on simply measuring the reduction in virus yield in the cell culture system, some mechanistic insights of emetine action have also been provided. For instance, in a study on human cytomegalovirus (HCMV), emetine was shown to target themouse double minute 2 homolog (MDM2)-p53 interaction, which is required for efficient HCMV replication (Mukhopadhyay et al., 2016). However, the precise natures of the cellular factors targeted by emetine and their roles in viral life cycle remain unknown. In this study, weprovide some novel mechanistic insights on the antiviral efficacy of emetine againstSARS-CoV-2, besides evaluating its protective efficacy against lethal challenge with infectious bronchitis virus (IBV, a chicken coronavirus) in embryonated chicken eggs.

## 2. Materials and Methods

### 2.1. Cells and viruses

African green monkey kidney (Vero)cells, available at the National Centre for Veterinary Type Cultures(NCVTC), Hisar were grown in Minimum Essential Medium (MEM) supplemented with 10% fetal bovine serum (FBS) (Sigma, St. Louis, USA) and antibiotics (Penicillin and Streptomycin). SARS-CoV-2 (Accession Number VTCCAVA294) was available in the national repository (NCVTC Hisar, Haryana, India). It was isolated in Vero cells from a human sample(naso-pharyngeal swab) which was received from Civil hospital Hisar, Haryana (India) to our facility at ICAR-National Research Centre on Equines (ICAR-NRCE), Hisar, Indiafor COVID-19 testing. SARS-CoV-2 was propagated in Vero cells in the Biosafety level 3 (BSL-3) laboratory of ICAR-NRCE, Hisar. The work was approved from the Institute Biosafety Committee and Review Committee on Genetic Manipulation (RCGM), Department of Biotechnology, Government of India (No. BT/BS/17/436/2011-PID). IBV was also available in the repository (NCVTC, Hisar) (Accession Number VTCCAVA168). It was propagated in specific pathogen free (SPF) embryonated chicken eggs which were procured from Indovax Pvt. Ltd., Hisar, India.

### 2.2. Inhibitors/Chemicals

Emetine[2*S*,3*R*,11*bS*)-2-[[(1*R*)-6,7-dimethoxy-1,2,3,4-tetrahydroisoquinolin-1-yl]methyl]-3-ethyl-9,10-dimethoxy-2,3,4,6,7,11*b*-hexahydro-1*H*-benzo[a]quinolizine]**(Fig. 1)**, FR180504 [ERK (extracellular signal-regulated kinase)] inhibitor, CGP57380 [MNK1 (MAPK-interacting kinase 1)] inhibitor, 4EGI-1[eIF4E(eukaryotic translation initiation factor 4E)] inhibitor and Apigenin [4′, 5, 7-Trihydroxyflavone, 5,7-Dihydroxy-2-(4-hydroxyphenyl)-4-benzopyrone] were procured from Sigma (Steinheim, Germany). Emetine was dissolved in phosphate buffered saline (PBS; vehicle control) whereas all other inhibitors were dissolved in DMSO. DMSO was used at a final concentration of 0.05%.3,3′-Diaminobenzidine (DAB) Enhanced Liquid Substrate System tetrahydrochloride and BCIP®/NBT solution (premixed) were procured from Sigma-Aldrich (St. Louis, USA).

**Fig. 1.**
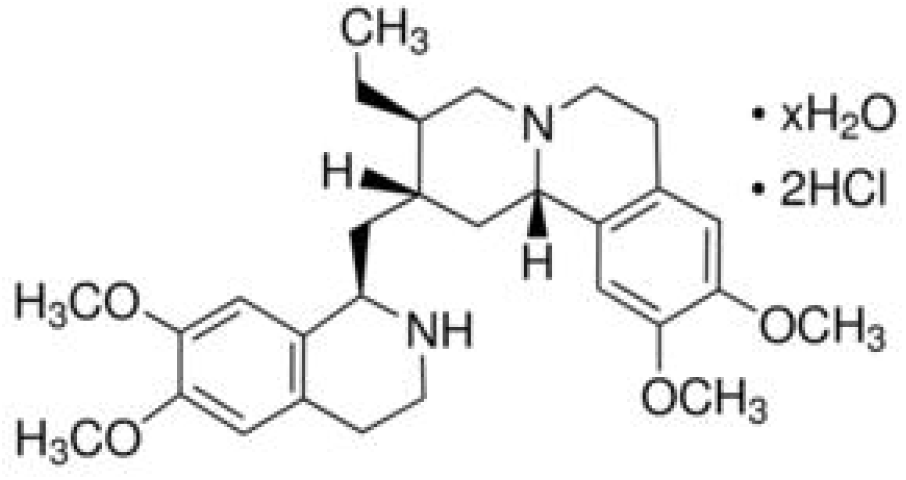
Structure of emetine.

### 2.3. Antibodies

eIF4E monoclonal antibody (5D11) and Phospho-eIF4E (Ser209) polyclonal antibodies were received from Invitrogen (South San Francisco, CA, USA). Mouse anti-GAPDH (Glyceraldehyde 3-phosphate dehydrogenase, house-keeping control protein) primary antibody, Anti-Mouse IgG (whole molecule)−Alkaline Phosphatase antibody (produced in goat) and Anti-Rabbit IgG (whole molecule)–Peroxidase antibody (produced in goat) were received from Sigma-Aldrich (St. Louis, USA). Rabbit anti-human IgG–HRP was procured from GeNei™, Peenya (Bangalore, India). Human serumfrom a COVID-19 confirmed patientwascollected at 2 weeks following recovery and obtained after due consent from the patient.

### 2.4. Determination of the cytotoxicity of emetine in Vero cells

The cytotoxicity (Kumar et al., 2008)of emetine in Vero cells was determined by MTT assay as described previously by our group(Khandelwal et al., 2017).

### 2.5. Determination of the lethal dose 50 (LD_50_) of emetine in embryonated chicken eggs

To determine LD_50_, 100 μl of 5-fold serial dilutions (5000-0.32 ng/egg) of emetine (5 eggs/dilution) or equal volumes of PBS (vehicle control)were injected in SPF embryonated chicken eggs via allantoic route. The eggs were observed daily for mortality of the embryos. The LD_50_ was determined by Reed-Muench method.

### 2.6. In ovo antiviral efficacy of emetine against IBV

SPF embryonated chicken eggs, in quadruplets, were inoculated with 5-fold serial dilutions of emetine (100-0.8 ng/egg in 100 μl PBS)via allantoicroute and subsequently infected with IBVategg infective dose 50 (EID_50_) of 100. The eggs were visualizeddaily for up to 6 days post-inoculation for mortality of the embryos. Effective concentration 50 (EC_50_) was determined by the Reed-Muench method.

### 2.7. Attachment assay

Vero cells, in triplicates were pre-incubated with 500 nM emetine or equivalent volume of PBS for 1 h and then infected with SARS-CoV-2 at MOI of 5 for 2 h at 4°C. The cells were then washed 5 times with PBS and the cell lysates were prepared by rapid freeze-thaw method. The viral titres were determined by plaque assay.

### 2.8. Entry assay

The effect of emetine on SARS-CoV-2 entry was assessed using a previously described assay (Khandelwal et al., 2014). Briefly, Vero cell monolayers (in triplicates) were pre-chilled to 4°C and infected with SARS-CoV-2 at MOI of 5 in the emetine-free medium for 1 h at 4°C to permit attachment. This was followed by washing and addition of fresh MEM containing the drug or its vehicle-control. Entry was allowed to proceed at 37°C for 1 h after which the cells were washed with PBS and incubated with MEMwithout any drug. The progeny virus particles released in the infected cell culture supernatant at 16 h were titrated by plaque assay.

### 2.9. Quantitative real-time PCR (RNA synthesis)

The levels of SARS-CoV-2 RNA in the infected cells were quantified by quantitative real-time PCR (qRT-PCR). Viral RNA was extracted by QIAamp Viral RNA Mini Kit (Qiagen, Hilden, Germany). cDNA was synthesized as per the protocol described by the manufacturer (Fermentas, Hanover, USA) using either oligo dT or random hexamer primers. qRT-PCR was carried out with a 25 μl reaction mixture containing gene specific primers, template and iTaq™ Universal SYBR® Green Supermix (Bio-Rad, USA) and run on QuantStudio™ 3 Real-Time PCR System (Thermo Fisher Scientific, Massachusetts, USA). Thermalcycler conditions were as follows: a denaturation step of 5 min at 95°C followed by 40 cycles of amplification (30 s at 95°C, 30 s at 59°C, and 30° s at 72°C). The levels of viral RNA, expressed as threshold cycle (CT) values, were analyzed to determine relative fold change in SARS-CoV-2 RNA copy numbers (E gene) as described previously (Kumar et al., 2016). The primers used for amplification of SARS-CoV-2 E gene were as follows: [forward primer: 5’-ACCGACGACGACTACTAGCG-3’and reverse primer: 5’-AGCTCTTCAACGGTAATAGTACCG-3’.

### 2.10. Virus release assay

Confluent monolayers of Vero cells, in triplicates, were infected with SARS-CoV-2 for 1 h at MOI of 5, followed by washing with PBS and addition of fresh MEM. At 10 hpi, cells were washed 5 times with chilled PBS followed by addition of fresh MEM containing 500nM emetine or PBS. Virus release at various time points (post-emetine addition) was quantified by plaque assay.

### 2.11. Western blot analysis

Confluent monolayers of Vero cells were infected with SARS-CoV-2 at MOI of 5for 1h, followed by washing with PBS and addition of fresh MEM. Cell lysates were prepared at 12 hpi in RIPA buffer (Kumar et al., 2011)and subjected to Western blot analysis to probe the viral/cellularproteins.

### 2.12. Chromatin immunoprecipitation (CHIP) assay

CHIP assay was carried out to evaluate the interaction of viral mRNA with a cellular cap-binding protein (eIF4E) as per the previously described method (Bier et al., 2011; Carey et al., 2009) with some modifications. Briefly, Vero cells, in triplicates were infected with SARS-CoV-2 at MOI of 5. At 10hpi, when the RNA levels were expected to be at its peak, the cells were treated with 1% formaldehyde for 10 min to covalently cross-link interacting proteins and nucleic acid. Thereafter, the cross-linking reaction was stopped by addition of 125 mM glycine (final concentration) followed by washing the cells with ice-cold PBS. The cells lysates were prepared in immunoprecipitation (IP) buffer [150 mM NaCl, 50 mM Tris-HCl (pH 7.5), 5 mM EDTA, 0.5% NP-40, 1% Triton X-100 plus protease and phosphatase inhibitor cocktail] and sonicated in a Qsonica Sonicator Q500 (Qsonica, Newtown, CT, USA)(6 pulse of 15 sec at amplitude of 40%). The cell lysates were then centrifuged for 10 min at 12,000g. The clarified cell lysateswere mixed with 10 units of RiboLock RNase Inhibitor (Thermo Scientific, USA) and then incubated with the phospho-eIF4E antibody (reactive antibody),phospho ERK antibody (nonreactive antibody) or equivalent volume of IP buffer (beads control) for 45 minutes at room temperature. Thereafter,40μl (5 ng/μl) of Protein A Sepharose® slurry, prepared as per the instruction of manufacturer (Abcam, USA) was added into each reaction and incubated overnight at 4°C on a rotary platform. The beads were then washed 5 times in IP buffer (without protease inhibitors). To reverse the cross-linking, the complexes were then incubated withProteinase K (20mg/ml final concentration) at 56°C for 40 minutes. Finally, the reaction mixtures were centrifuged at 12,000g for 1 min and the supernatant was subjected to cDNA preparation and quantitation of SARS-CoV-2 RNA (E gene) by qRT-PCR.

## 3. Results

### 3.1. In vitro antiviral efficacy of emetine against SARS-CoV-2

Emetine at concentrations of ~500 nMhad no effects onthe viability ofVero cells (**Fig. 2a**), though at higher concentrations (≥2000 nM), it was found to be quite toxic to the cells (**Fig. 2a**). The CC_50_ was determined to be 1603.8 nM. To analysethe effect ofemetine on cell free virions (virucidal effect), infectious virions were incubated with indicated concentrations of emetine which wasfollowed by determination of theresidual infectivityof SARS-CoV-2 virions by plaque assay. As shown in **Fig. 2b,** viral titres were comparable in the emetine and vehicle control-treated groups which suggestedthat emetine has no virucidal effect on the cell-free SARS-CoV-2 virions. To determine the*in vitro*antiviral efficacy of emetine, the yields of infectious SARS-CoV-2 virionswere measured in the presence of various non-cytotoxic concentrations of emetineor vehicle-control.Emetine was found to decrease the SARS-CoV-2 replication **(Fig. 2c)** in a dose-dependent manner. The EC_50_ was determined to be 0.147 nM. The selective index (CC_50_/EC_50_) was calculated to be 10910.4.

**Fig. 2.**
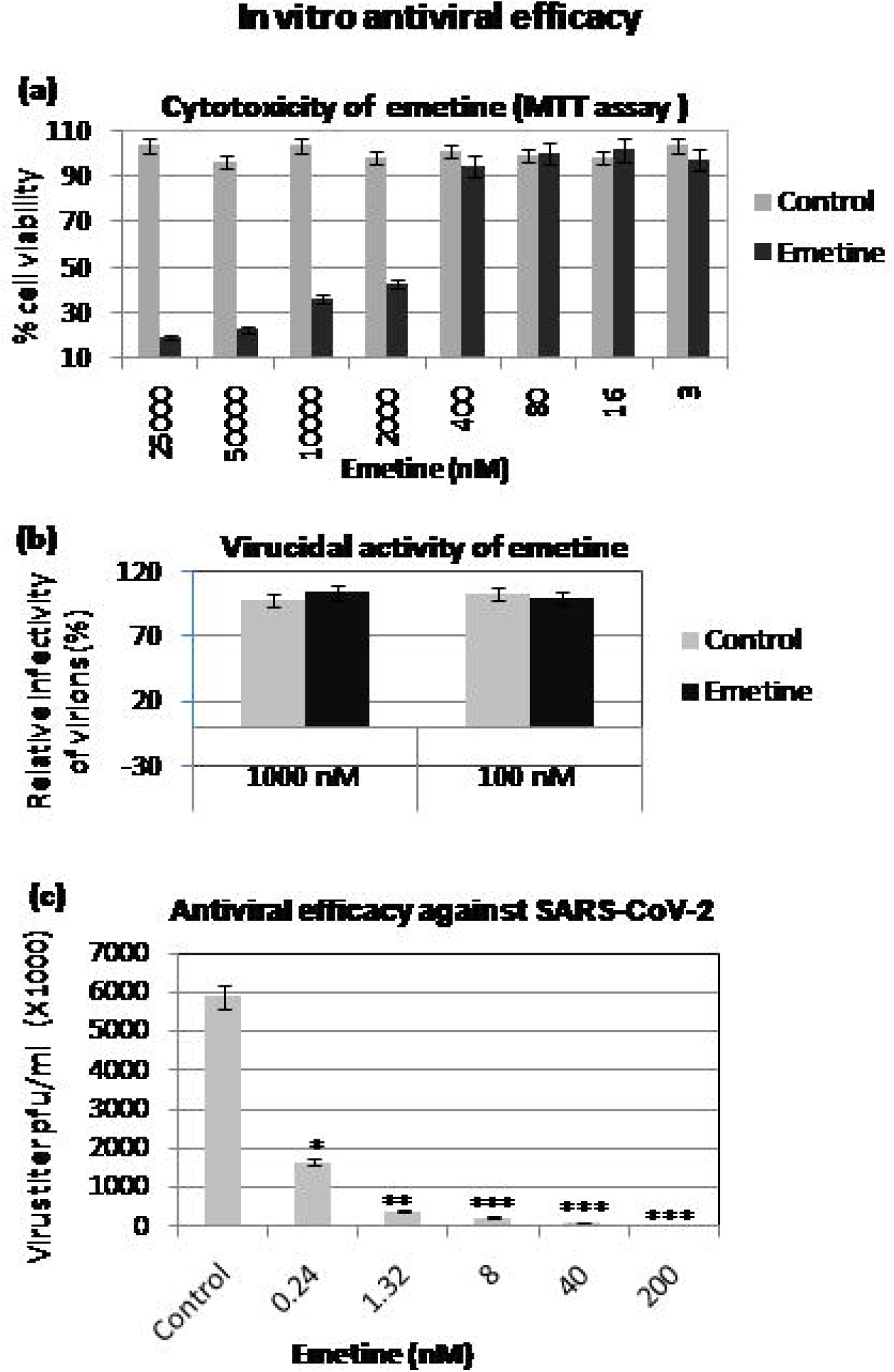
*In vitro* antiviral efficacy of emetine against SARS-CoV-2. ***(a)Determination of the cytotoxicity(MTT assay):*** Indicated concentrations of emetine or equivalent volumes of vehicle control (PBS) were incubated with cultured Vero cells for 96 h and % cell viability was determined by MTT assay. ***(b)Virucidal activity:*** Indicated concentrations of the emetine or equivalent volumes of PBS were mixed with 10^6^ pfuof SARS-CoV-2 and incubated for 90 min at 37°C after which residual viral infectivity (% of control) was determined by plaque assay. ***(c)In vitroantiviral efficacy:*** Vero cells, in triplicates, were infected with SARS-CoV-2 atMOI of 0.1 in the presence of indicted concentrations of emetine or vehicle-control. The virus particles released in the infected cell culture supernatants at 24 hpi were quantified by plaque assay. Values are means ± SD and representative of the result of at least 3 independent experiments. Pair-wise statistical comparisons were performed using Student’s t test(* = P<0.05, (** = P<0.01, (*** = P<0.001).

### 3.2. In ovo antiviral efficacy of emetine against IBV

Anti-coronavirus potential of emetine was further evaluated *in ovo*by virulent IBV challenge in an embryonated chicken egg infection model. Initially, we determined the lethal dose of emetine by inoculating SPF embryonated chicken eggs with serial dilutions of emetine. As shown in **Fig. 3a**, mortality of the embryo was observed at emetine concentration ≥200 ng/egg but not at the lower concentrations. The LD_50_ was determined to be 365.7 ng/egg.

**Fig. 3.**
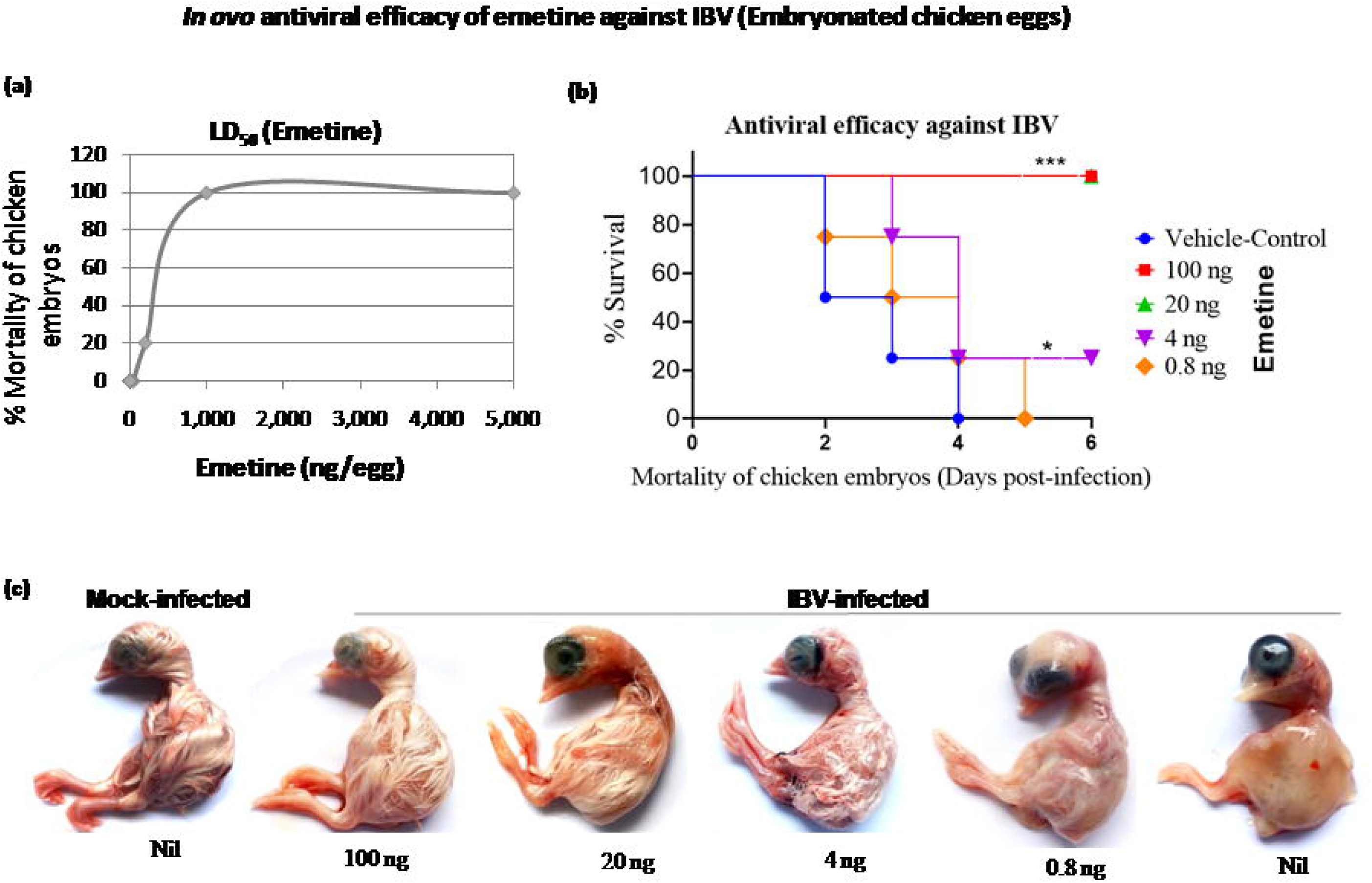
*In ovo* antiviral efficacy of emetine against IBV. **(a) *Determination of LD_50_***: SPF embryonated chicken eggs were injected with 5-fold serial dilutions of emetine (5 eggs/dilution) or vehicle control via allantoic route and observed daily for mortality of the embryos for up to 5 days. LD_50_ was determination by the Reed-Muench method. ***(b**)**In ovo antiviral efficacy:*** SPF embryonated chicken eggs were infected with IBV at EID_50_of 100 via allantoic route in the presence of indicated concentrations of emetine or PBSand observed daily for mortality of the embryos. Duration of the survival of chicken embryos following IBV challenge as determined by Kaplan-Meier (survival) curve is shown. LD_50_ was determined by the Reed-Muench method. Statistical comparisons in survival curves were made using Log-rank (Mantel-Cox) Test using GraphPad Prism 7.02. ***(c)*** Morphological changes in the chicken embryos at different drug regimens following IBV challenge is shown. *= P<0.05, ***= P<0.001.

For evaluation of *in ovo*anti-IBV efficacy of emetine, eggs were infected with IBV along with the indicated concentrations of emetine. As shown in **Fig. 3b,** emetine prevented the chicken embryos against lethal IBV challenge in a dose-dependentmanner. **N**o mortalities were observed at emetine concentration of ≥20 ng whereas a partial (significant) protection was obtained up to 4 ng. Besides, emetine administration also resulted in normal development of embryos as compared to vehicle-control-treated group wherein stunted growth and defective feather development was observed **(Fig. 3c).** At an EC_50_ of 2.3 ng/egg, the therapeutic index (LD_50_/EC_50_) of emetine was determined to be 159.1. Taken together, we conclude that emetine prevents the chicken embryos against lethal challenge with IBV.

### 3.3. Emetinedoes not affect attachment, entry and release of SARS-CoV-2

In order to evaluate which specific step(s) of SARS-CoV-2 are affected by emetine, we first determined the life cycle of SARS-CoV-2in one-step growth curve wherein Vero cells were infected with high multiplicity of infection (MOI=5) followed by quantification of the virus released in the infected cell culture supernatant at different times post-infection. As shown in **Fig. 4a,** the virus titres were comparable at 1 hpi, 2 hpi or 4 hpi. However, a sharp rise in the viral titres was observed at 8 hpi. This was presumably due to the production of new progeny virus particles and hencesuggestive of the completion of viral life cycle. After 8 hpi viz; 10 hpi and 12 hpi, the virus entered intothe stationary phase**(Fig. 4a)**. Keeping these results in view and in agreement with others (Keyaerts et al., 2005), the life cycle of SARS-CoV-2 was determined to be 8-10 h in Vero cells.

**Fig.4.**
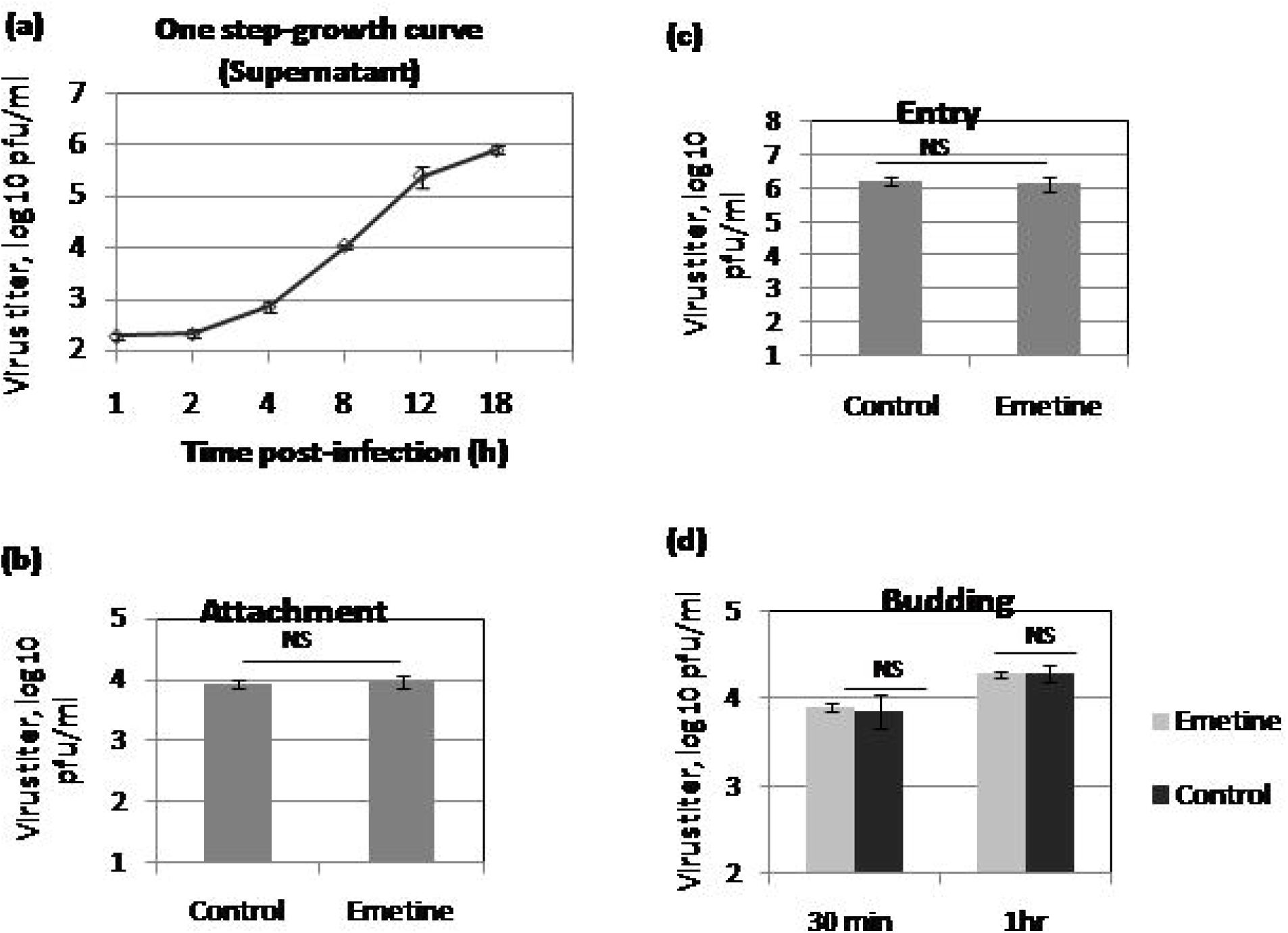
Effect of emetine on specific steps of viral life cycle. ***(a)One-step growth curve***:Confluent monolayers of Vero cells, in triplicates,were infected with SARS-CoV-2 at MOI of 5, and thereafter washed with PBS and fresh MEM was added. Infectious progeny virus particles released in the cell culture supernatant at indicated time points were quantified by plaque assay. ***(b):Attachment assay:*** Vero cells were pre-incubated with 500 nM emetine or vehicle-control for 1 h and then infected with SARS-CoV-2 at MOI of 5 for 2 h at 4°C. The cells were then washed 5 times with PBS and the cell lysates were prepared by rapid freeze-thaw method. The virus titres were determined by plaque assay. ***(d)Viral entry:*** Vero cell monolayers, in triplicates, were pre-chilled to 4°C and infected with SARS-CoV-2 at MOI of 5 in emetine-free medium for 1h at 4°C to permit attachment. This was followed by washing with PBS and addition of fresh MEM containing 500 nM emetine or vehicle-control. Entry was allowed to proceed at 37°C for 1h after which the cells were washed again with PBS to remove any extracellular viruses and incubated with cell culture medium without any inhibitor. The progeny virus particles released in the infected cell culture supernatants at 16 h in the treated and untreated cells were titrated by plaque assay. ***(e)Virus release assay***: Confluent monolayers of Vero cells, in triplicates, were infected with SARS-CoV-2, for 2 hat MOI of 5 followed by washing with PBS and addition of fresh MEM. At 10 hpi, cells were washed 6 times with chilled PBS followed by addition of fresh MEM containing 500 nM emetine or vehicle-control. Virus release at indicated time points (post-emetine addition) was quantified by plaque assay. Values are means ± SD and representative of the result of at least 3 independent experiments. NS represents no statistical significance.

Typically, the receptor-virus interactions are specific and the binding is independent of energy or temperature. The attachmentassay was carried out at 4°C which allowed only attachment but restricted other post-attachment steps of the viral life cycle which are energy and temperature dependent. As shown in **(Fig. 4b),** we did not observe any significant difference in the viral tittersattached in the presence of emetine or vehicle-control. In order to determine whetherthe pre-attached virus was able to enter the cells in the presence of emetine, a standard entry assay was also performed. Emetine was not shown to affect viral entry as the viral titres were comparable between emetine-treated and vehicle-control-treated cells **(Fig. 4c).** To determine the effect of emetine on virus release (budding), it was applied at the time when the virus was presumed to be completing its life cycle viz; ~10 hpi. As shown in **Fig. 4d,** no significant difference in the viral titres was observed between emetine-treated and control-treated cells, suggesting that emetine does not affect SARS-CoV-2 release from the infected cells.

### 3.4. Emetine treatment reduces SARS-CoV-2 RNA and proteins synthesis

In order to determinethe effect of emetine on the synthesis of viral genome, we first evaluated the kinetics of the viral RNA synthesis in the infected cells (cell pellet). As shown in **Fig. 5a,** the amount of viral RNA in the infected cell pellet was comparable at 1 hpi, 2 hpi, 4 hpi and 6 hpi. The detectable amounts of viral RNA at these early time points represent the virus particles which entered in the target cells following infection (input). A sharp rise in the viral RNA was observed at 8 hpi which peaked at 10 hpi before declining sharply at 12 hpi **(Fig. 5a).**

**Fig.5.**
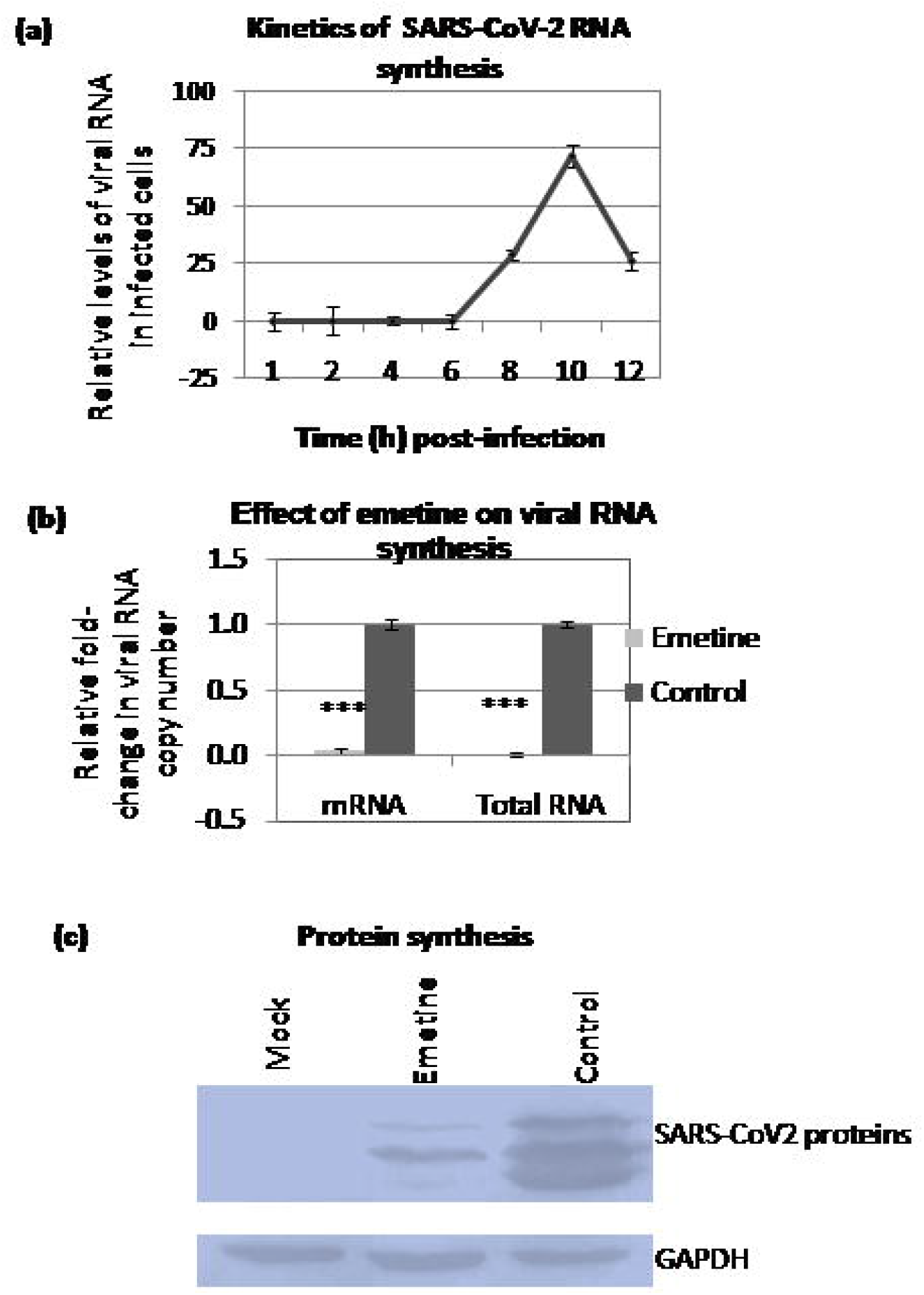
Emetine treatment results in decreasedsynthesis of SARS-CoV-2 RNA and proteins. ***(a):Kinetics of SARS-CoV-2 RNA synthesis:*** Confluent monolayers of Vero cells, in triplicates, were infected with SARS-CoV-2 at MOI of 5, followed by washing with PBS and addition of fresh MEM. Cells were scrapped at indicated time points and subjected for the quantitation of viral RNA by qRT-PCR. cT values were normalized with β-actin house-keeping control gene and relative fold-change was calculated by ΔΔ Ct method. ***(b)Effect of emetine onRNA synthesis:*** Confluent monolayers of Vero cells, in triplicates, were infected with SARS-CoV-2 for 1 h at MOI of 5. Emetine was added at 4 hpi and cells were harvestedat 10 hpi to determine the levels of SARS-CoV-2 RNA by qRT-PCR. Threshold cycle (Ct) values were analysed to determine relative fold-changein copy numbers oftotal RNA and mRNA. Values are means ± SD and representative of the result of at least 3 independent experiments. Pair-wise statistical comparisons were performed using Student’s t test (* = P<0.05,*** = P<0.001). ***(c)Protein synthesis:***Vero cells were infected with SARS-CoV-2 at MOI of 5, followed by washing with PBS and addition of fresh MEM. Cell lysates were prepared at 12 hpi to detect the viral proteins by Western blot analysis by using serum derived from a COVID-19 positive patient. The levels of viral proteins **(upper panel)**, along with housekeeping GAPDH protein (**lower panels**) are shown.

To evaluate the effect of emetine on synthesis of SARS-CoV-2 RNA, emetine was applied when early steps of the virus life cycle (attachment/entry) were expected to occur (~4hpi) and the cells were harvested when the levels of viral RNA were expected to be maximum (~10 hpi, **Fig. 5a).** As shown in **Fig. 5b**, a highly significantreduction in the relative copy numbers of viral RNAas well as viral mRNA was observed in emetine-treated cells whichsuggestedthat emetine could affect viral genome synthesis in infected cells.

Emetine-induced decreasedsynthesis of viral genome could be due to the reduced synthesis of viral proteins (viral polymerase and other accessory proteins required for virus replication).Nevertheless, Western blot analysis of cells infected withSARS-CoV-2 (**Fig. 5c, upper panel**) showed decreased levels of viral proteins in the presence of emetine. However, there were similar levels of cellular housekeeping proteinGAPDH in emetine-treated and untreated cells (**Fig. 5c, lowerpanel**)which suggested that emetine does not lead to a general shut off /inhibitionof cellular protein synthesis(at least at the concentration we employed).

### 3.5. Emetine inhibits interaction of SARS-CoV-2 mRNA and cellular eIF4E

We further explored the possible inhibitory mechanism of emetine in suppressingthe synthesis of SARS-CoV-2 proteins.In coronaviruses, viral mRNA translation takes place in a cap-dependent manner wherein the eIF4E plays a central role in the initiation of translation(Cencic et al., 2011; Gordon et al., 2020; Kumar et al., 2018; Müller et al., 2018). Upon activation (phosphorylation) by upstream kinase(s), elF4E binds to 5′ cap of mRNA to initiate translation (Kumar et al., 2018). We first evaluated the impact of the inhibition of eIF4E activity on SARS-CoV-2 replication. As shown in **Fig. 6a,** at a non-cytotoxic concentration [(0.5 μg/ml, determined previously by our group (Khandelwal et al., 2020)], addition of 4EGI-1 resulted in a highly significant reduction invirus yield**(Fig. 6a)** whichsuggested that eIF4E is essential for SARS-CoV-2 replication.

**Fig. 6.**
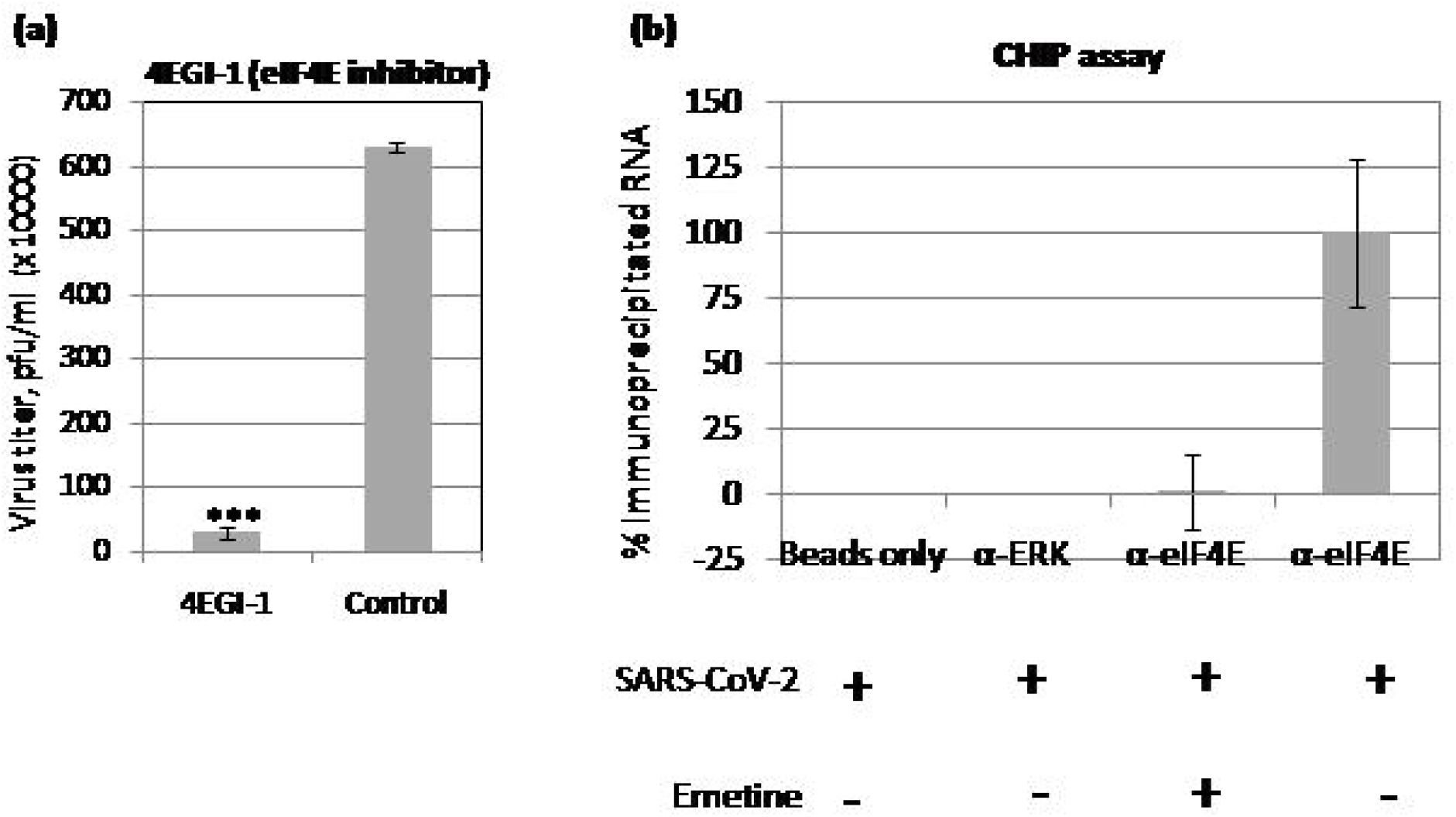
Emetine inhibits interaction of SARS-CoV-2 mRNA and eIF4E. ***(a):eIF4E is required for efficient SARS-CoV-2 replication.*** Vero cells, in triplicates, were infected with SARS-CoV-2 at MOI of 0.1 in the presence of 0.5 μg/ml of 4EGI-1 (eIF4E inhibitor) or equivalent volume of DMSO (vehicle-control). The virus yield in the infected cell culture supernatantwas quantified by plaque assay. ***(b): CHIP assay:*** Vero cells, in triplicates were infected with SARS-CoV-2 at MOI of 5 followed by washing with PBS and addition of fresh MEM containing emetine or vehicle control(s). At 10 hpi, cell lysates were prepared as per the procedure described for CHIP assay (materials and method section). The clarified cell lysates were incubated with α-eIF4E (reactive antibody), α-ERK (nonreactive antibody) or equivalent volume of IP buffer (Beads control) followed by incubation with Protein A Sepharose® slurry. The beads were then washed five times in IP buffer. To reverse the cross-linking, the complexes were then incubated with Proteinase K. Finally, the reaction mixtures were centrifuged and the supernatant was subjected to cDNA preparation and quantitation of SARS-CoV-2 RNA (E gene) by qRT-PCR.Values are means ± SD and representative of the result of at least 3 independent experiments. Pair-wise statistical comparisons were performed using Student’s t test (* = P<0.05, (** = P<0.01, (*** = P<0.001).

Next we evaluated if emetine inhibits interaction of the viral RNAwith eIF4E (cap-binding protein). At 10 hpi when SARS-CoV-2 RNA was expected to be at its peak level, cells were covalently cross-linked and evaluated for viral RNA and eIF4E interaction in a CHIP assay. As shown in **Fig. 6b,** α-eIF4E (reactive antibody) but not α-ERK (non-reactive antibody) or beads control immunoprecipitated SARS-CoV-2 RNA. The levels of viral RNA immunoprecipitated by α-eIF4E were >99.9% lower in emetine-treated cells as compared to the PBS-treated cells (**Fig. 6b)** which confirmed that emetine inhibits eIF4E/SARS-CoV-2 mRNA interaction. In qRT-PCR, the levels of SARS-CoV-2 RNA in α-ERK-treated (but notα-eIF4E-treated cell aliquots) were undetectable (Ct values undetermined) which suggested that α-eIF4E specifically interacted with SARS-CoV-2 RNA(**Fig. 6b**).

### 3.6. ERK/MNK1/eIF4E cell signalling pathway is prerequisite for SARS-CoV-2 replication

eIF4E is activated via RTK/ERK/MNK1 signalling axis (Kumar et al., 2018).As shown in **Fig. 7,** in addition to the inhibitory effect of eIF4E inhibitor, at a non-cytotoxic concentration, addition of the inhibitors targetingupstream eIF4E kinases such asMNK1 (CGP57380: 0.5 μg/ml) and ERK (FR180204: 0.2 μg/ml) also resulted in decreased SARS-CoV-2 replication suggesting that the ERK/MNK1/eIF4E signalling is prerequisite for SARS-CoV-2 replication. The CC_50_ values of eIF4E, Apigenin, CGP57380 and FR180204 in Vero cells has been described previously by our groups (Khandelwal et al., 2020). Apigenin, a dietary flavanoid was previously shown to decrease eIF4E phosphorylation in buffalopox virus (BPXV) infected Vero cells(Khandelwal et al., 2020).Addition of a non-cytotoxic concentration (2.5μg/ml) of Apigenin resulted in reduced production ofSARS-CoV-2 in Vero cells **(Fig. 7)** which further confirmed the virus supportive role of eIF4E in SARS-CoV-2 replication.

**Fig. 7.**
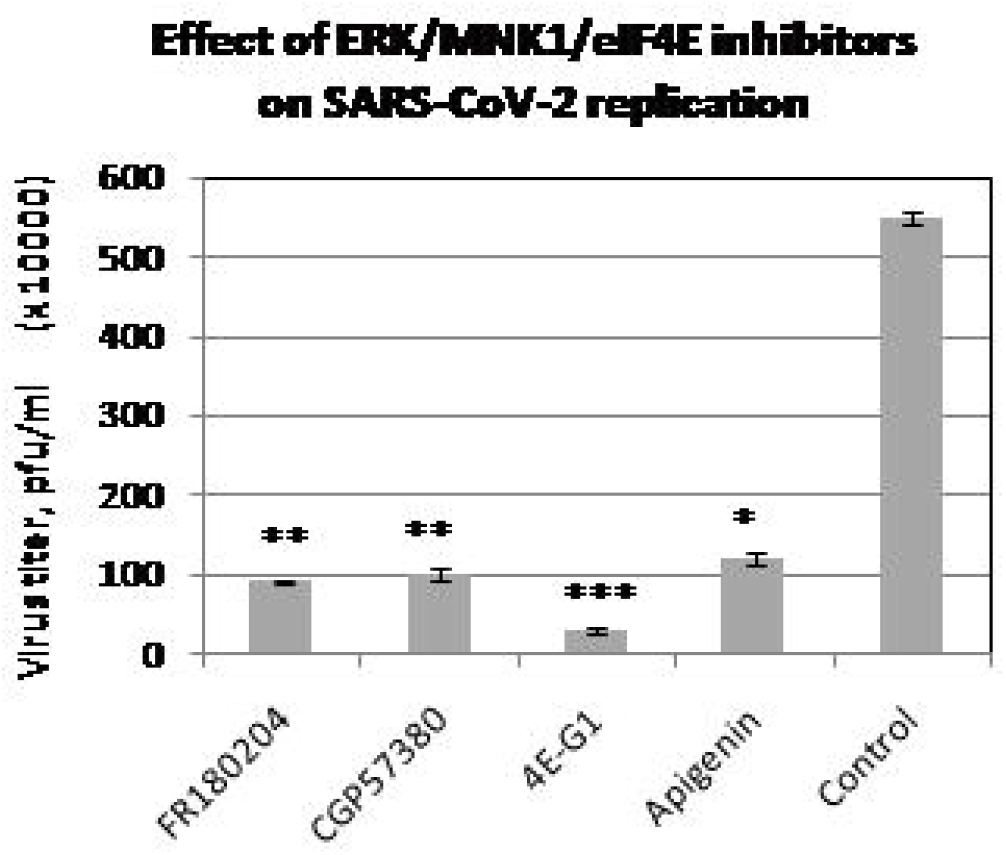
ERK/MNK1/eIF4E signalling is prerequisite for SARS-CoV-2 replication. Vero cells, in triplicates, were infected with SARS-CoV-2 in the presence of the indicated inhibitors viz; ERK inhibitor (FR180204: 0.2 μg/ml), MNK1 inhibitor (CGP57380: 0.5 μg/ml), eIF4E inhibitor (0.5 μg/ml) and Apigenin (2.5 μg/ml).The virus yield in the infected cell culture supernatants at 24 hpi was quantified by plaque assay.

## 4. Discussion

We along with some other groups have demonstrated the*in vitro* antiviral efficacy of emetine against some RNA and DNA viruses (Chaves Valadao et al., 2015; Choy et al., 2020; Deng et al., 2007; Khandelwal et al., 2017; Mukhopadhyay et al., 2016; Ramabhadran and Thach, 1980). While the development of entirely new antiviral drugs may take several months or even years, repurposing the existing approved drugs could save time and a lot of investments. Following the emergence of COVID-19, we evaluated the*in vitro* antiviral activity of emetine against SARS-CoV-2. In this study, we have provided some novel mechanistic insights on the antiviral efficacy of emetine against SARS-CoV-2, besides analysingthe antiviral potential of emetine againsta lethal challenge with IBV for the first time.

Emetine inhibits protein synthesis in mammalian cells. A non-cytotoxic concentration of emetine is also known to inhibit the replication of certainviruses. While most of the studies on the inhibitory effect of emetine on virus replication are simply based on measuring the virus yields in cell culture(Choy et al., 2020; Ramabhadran and Thach, 1980; Yang et al., 2018),somemechanistic insights have also been provided. Emetine can directly inhibit viral polymerase(Chaves Valadao et al., 2015; Khandelwal et al., 2017; Yang et al., 2018), although the major antiviral activity of emetine is believed to be mediated via targeting certain cellular factors (Khandelwal et al., 2017). Depending on the nature of the virus involved, emetine could target different step(s) of virus replication cycle. For instance, in Zika virus (ZIKV), emetine was shown to inhibit NS5 polymerase activity, besides inhibiting viral entry (mediated via disruption of lysosomal functions)(Yang et al., 2018). In human cytomegalovirus (HCMV) infection, emetine was shown to inhibit HCMV replication after entry but before initiation of DNA synthesis(Mukhopadhyay et al., 2016). Likewise, in rabies virus, emetine was also shown to block the axonal transport of the virus particles (MacGibeny et al., 2018). Similarly, in vaccinia virus, it was shown to interfere with the virus assembly (Deng et al., 2007). Most of these studies provide insights on the inhibitory effect of emetine on the specific step(s) of viral life cycle. However, the precise molecular mechanism of emetine action remains unknown.

Viruses are well known to exploit several cellular factors to effectively transcribe and translate their genome(Kumar et al., 2020). Like several other viruses, coronaviruses also synthesizetheir protein in cap-dependent manner wherein eIF4E plays a critical role in the initiation of translation(Nakagawa et al., 2016). Since emetine was found to decrease synthesis of viral proteins, we hypothesize ifinhibitory effect of emetine is mediated via disrupting viral mRNA and eIF4E interaction. As in the case of other coronaviruses (Nakagawa et al., 2016), inhibiting the cellular eIF4E activityresulted in decreasedSARS-CoV-2 replication which suggested the virus supportive role of eIF4E.Further, in a CHIP assay, we demonstrated that emetine inhibitsinteraction of the viral RNA with cap-binding protein eIF4E**(Fig. 6b** and**Fig. 8).** eIF4E activation is commonly mediated via RTK/ERK/MNK1/eIF4Esignalling axis(Kumar et al., 2018).Blocking these upstream eIF4E kinases resulted in reduced SARS-CoV-2 replication which suggested that ERK/MNK1/eIF4E signalling is prerequisite for SARS-CoV-2 replication**(Fig. 8).** Nevertheless previous studies suggest that coronaviruses induces eIF4E activation which plays a virus supportive role in coronavirus (including SARS-CoV-2)life cycle (Cencic et al., 2011; Gordon et al., 2020; Mizutani et al., 2004; Müller et al., 2018).

**Fig. 8.**
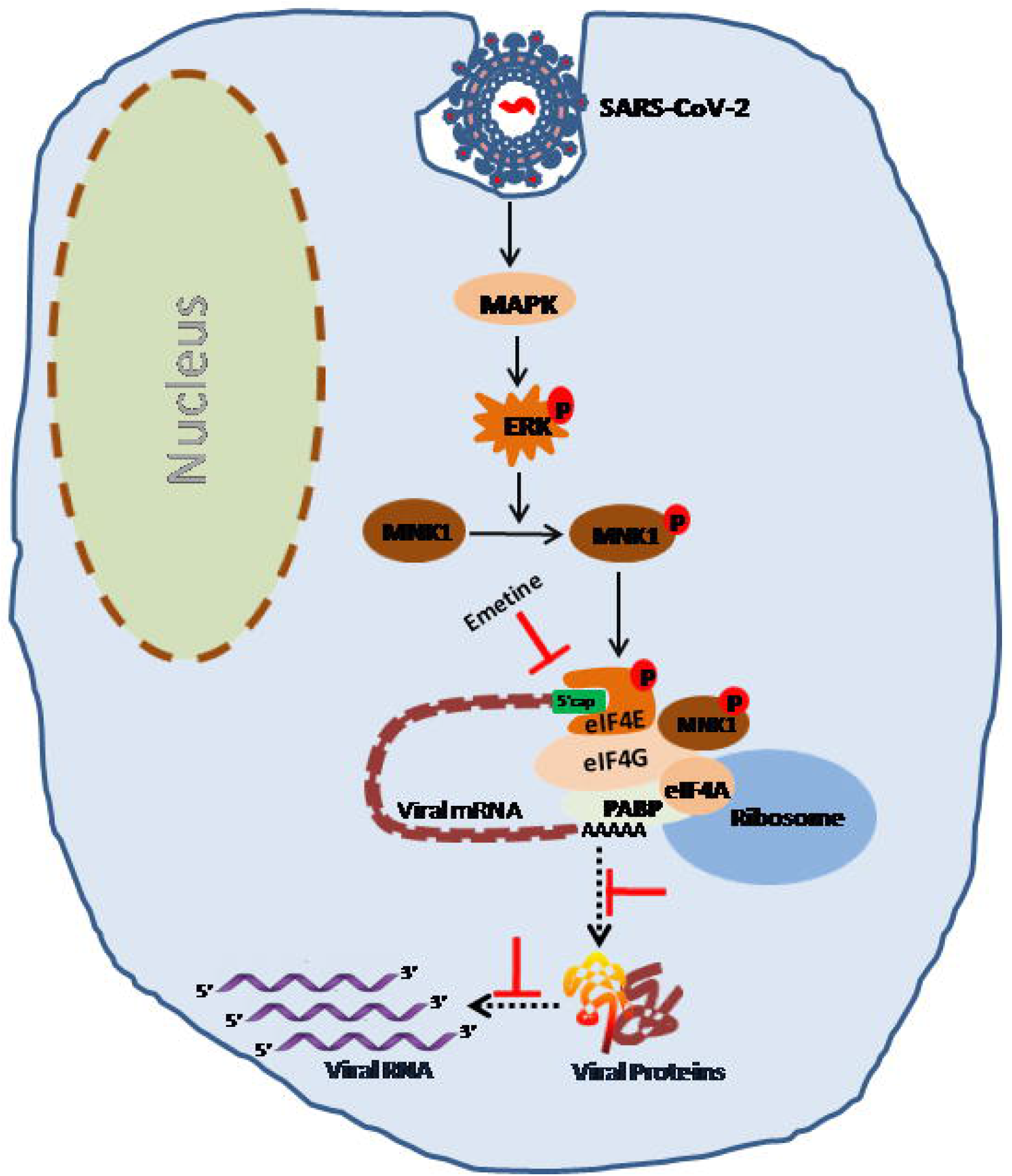
Schematic diagram ofthe possible inhibitory mechanism of emetine on SARS-CoV-2 replication. SARS-CoV-2exploitsMAPK/ERK/MNK1/eIF4Ecell signalling pathway to effectively replicate in the target cells. Activated eIF4E binds with 5’-cap structure of viral mRNA to initiates translation of viral proteins. Emetine blocks SARS-CoV-2 protein synthesis by inhibiting interaction of viralmRNA with eIF4E.

Since virus replication requires the presence of the viral structural and non-structuralproteins, their (viral polymerase), their restricted availability (in the presence of emetine) couldeventually block viral genome replicationwhich we have observed in our study in the form of reduced levels of viral RNA and mRNA. Alternatively, emetine may directly inhibit certain viral polymerases (Aguiar et al., 2007; Chaves Valadao et al., 2015). However, in a cell free virion polymerase assay, we could not observe any direct effect of emetine on the function of viral polymerase (data not shown) which suggests that the major mechanism of action of emetine is mediated via targeting the cellular factor(s), one (eIF4E) which is described in this study. Like eIF4E, emetine may also target other host-dependency factors. Additional studies that involve transcriptome/proteome/phospho-proteome/lipidome analysis in emetine-treated and-untreated cells are required to precisely elucidate the molecular mechanism of the action of emetine.

The high selective index of emetine suggests its potential as anti-SARS-CoV-2 agent. However, since emetine also targets cellular factors, *in vivo* cytotoxicity may be a potential difficulty of its use. For the treatment of amebiasis, emetine is given at 1 mg/kg body weight daily for up to 10 days without any side effects (Mastrangelo et al., 1973). As an anti-cancer agent (clinical trials), emetine was shown to be well tolerated when delivered intravenously at 1.5 mg/kg doses twice a week (Panettiere and Coltman, 1971). In a mouse CMV (MCMV) model, emetine inhibited virus replication at an oral dose of 0.1 mg/kg body weight(Mukhopadhyay et al., 2016)which is somewhat in the similar range required to protect chicken embryos against virulent IBVchallenge in this study (considering an embryo weight of 20 gram and an EC_50_ of 2.3 ng/egg). Although the route of administration and potential cumulative cytotoxicity needs to be determined in the case of COVID-19, the doses required for virus inhibition are substantially lower than the traditional emetine doses used to treat amebiasis and other ailments. Considering these facts in view, it is apparent that emetine has the potential to treat COVID-19.

*In vitro* antiviral efficacy against SARS-CoV-2 and its ability to protect chicken embryos against virulent IBV suggests that emetine could be repurposed to treat COVID-19. In addition, we provide a novel mechanistic insight into the antiviral activity of emetine. Emetine targets SARS-CoV-2 protein synthesis which is mediated by inhibiting the interaction of SARS-CoV-2 mRNA with the cellular cap-binding protein eIF4E.

## Acknowledgements

This work was supported by the Science and Engineering Research Board, Department of Science and Technology, Government of India [grant number CVD/2020/000103].

